# Cardiac metabolite exchange measured in pigs during myocardial ischemia and reperfusion

**DOI:** 10.1101/2025.08.04.668556

**Authors:** Ian Tamburini, Aleksandra Stamenkovic, Hosung Bae, Wonsuk Choi, Marcus Seldin, Cholsoon Jang, Amir Ravandi

## Abstract

Myocardial ischemia and reperfusion (I/R) injury are the primary contributors to death in patients with cardiovascular disease. While decades of research have elucidated the molecular players and biochemical mechanisms underlying I/R injury, how these pathologies influence the metabolic activities of the heart remains incompletely understood. Such a knowledge gap hampers the development of therapies aimed at mitigating the metabolic stresses of the heart during injury. Using comprehensive arteriovenous metabolomics in a highly relevant porcine I/R model, we report the metabolic landscape of cardiac metabolic changes after ischemia and during the reperfusion time course. Paradoxically, ischemia increases the cardiac uptake of circulating fatty acids while reperfusion for 60 minutes reverses this activity. By 120 minutes of reperfusion, the hearts resume the uptake of fatty acids, suggesting restoration of their metabolism. On the other hand, we found a strong release of amino acids by the heart only after 60-minute reperfusion, but not after 120-minute reperfusion, implicating I/R-induced transient protein degradation. In addition to these findings, we identified several previously unrecognized changes in cardiac metabolic inputs and outputs during I/R, including nucleotides, TCA cycle intermediates and creatine/creatinine. These data highlight the dynamic alterations in cardiac metabolism in response to I/R, providing insights into how to mitigate myocardial I/R injury.

## INTRODUCTION

Cardiovascular disease remains the leading cause of morbidity and mortality worldwide, with 85% of deaths due to myocardial infarction (MI) and stroke^1^. Following an acute MI, rapid restoration of blood flow either via percutaneous coronary intervention (PCI) or administration of thrombolytics is the first line of therapy. However, these treatments cause further damage known as ischemia/reperfusion (IR) injury^2^. During ischemia, the interruption of coronary blood flow leads to a profound disruption of myocardial metabolism, characterized by ATP depletion, intracellular acidosis, and accumulation of anaerobic metabolites such as lactate. These metabolic disturbances impair ion homeostasis, disrupt membrane integrity, and initiate cell death pathways. Reperfusion, although essential for salvaging viable myocardium, paradoxically exacerbates injury through a burst of reactive oxygen species (ROS), calcium overload, mitochondrial dysfunction, and the activation of inflammatory cascades^2–5^. This paradoxical damage contributes to infarct expansion, adverse cardiac remodeling, and long-term heart failure.

Metabolomics, the comprehensive profiling of small molecules, provides a powerful lens through which to examine the dynamic biochemical shifts that occur during I/R. By capturing changes in energy metabolism, oxidative stress, amino acid catabolism, and lipid signaling, metabolomics enables a systems-level understanding of tissue injury and repair mechanisms^6,7^. However, most I/R metabolomic studies rely on measuring snapshot metabolite levels in systemic blood or heart tissues, limiting insight into the metabolic fluxes or activities of the heart.

Arteriovenous (AV) metabolomics analysis, comparing metabolite concentrations between paired arterial and venous blood draining a specific tissue, provides a unique approach to quantify net metabolite uptake and release by the target tissue, which infers tissue-specific metabolic fluxes. This method has been successfully applied to investigate the metabolism of critical organs, including muscle, liver, adipose tissue, and brain, under various physiological and pathological conditions^8–15^.

Despite its potential, AV metabolomics has not been utilized to understand the metabolic flux changes in the heart during I/R. In this study, we employed paired arterial and venous sampling and comparative metabolomics in a porcine model of myocardial I/R, illuminating several distinct patterns of metabolic changes of the heart during I/R. In particular, we observed dynamic changes of fatty acid and amino acid uptake and release by the heart during ischemia followed by reperfusion time-course.

## RESULTS

### Ischemia and reperfusion distinctively influence systemic circulating metabolome

To examine how I/R affects metabolism at the whole-body level versus in the heart, we performed untargeted metabolomics on systemic arterial blood (sampled from the femoral artery) and draining venous blood from the heart (sampled from the coronary sinus). Using liquid chromatography-tandem mass spectrometry (LC-MS), we measured a total of 464 circulating metabolites, whose chemical identities were confirmed by authentic standards. To capture the dynamic metabolic changes during I/R, we compared metabolite levels across four time points: baseline (no I/R), 60 minutes after ischemia (before reperfusion), and 60 minutes and 120 minutes after reperfusion (**Fig. 1A**).

**Figure 1.**
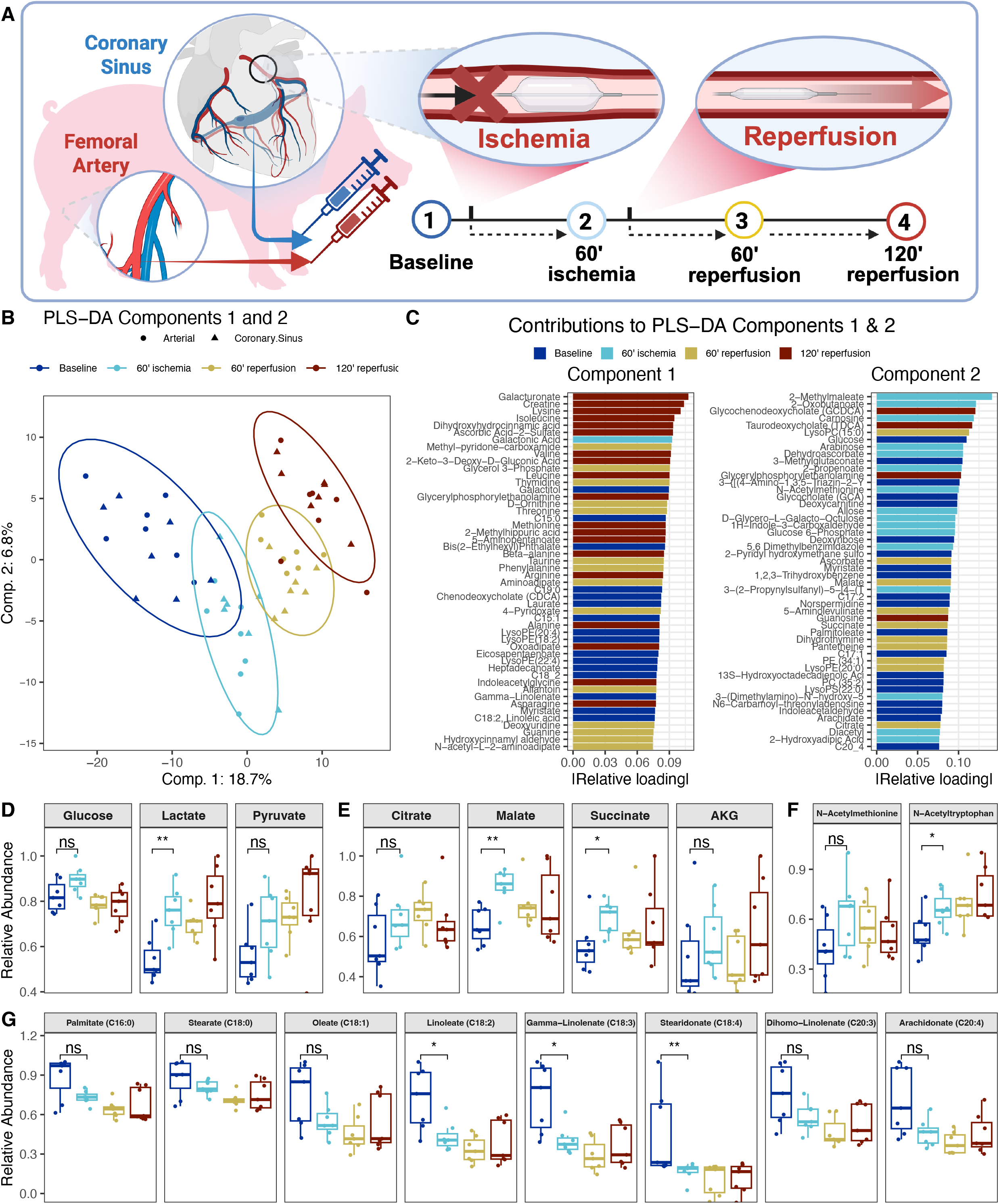
Comprehensive arteriovenous blood metabolomic profiling in the porcine model of ischemia and reperfusion time course. **A,** Schematic of the experimental workflow. Arterial blood from the femoral artery and venous blood from the coronary sinus were sampled during I/R. **B,** Partial Least Squares Discriminant Analysis (PLS-DA) of metabolomic profiles across different time points. **C,** Top metabolites contributing strongly to the components 1 and 2. **D-G:** Relative arterial abundance of the indicated metabolites in central carbon pathways at different time points. *p<0.05, **p<0.01 by Student’s t-test. ns, not significant.

We then performed partial least squares discriminant analysis (PLS-DA) using the mixOmics package (v6.22.0) in R^16^ to determine which patterns of variation in the metabolome most strongly distinguished the circulating metabolome during I/R. The analysis revealed apparent time-point-specific clustering of arterial and venous blood metabolome along components 1 and 2 (**Fig. 1B**). Top 50 metabolites that contribute to component 1 were mainly enriched in blood samples collected after reperfusion (red and brown in **Fig. 1C**), suggesting that component 1 reflects the metabolic changes driven by reperfusion. Intriguingly, the majority of these metabolites were amino acids (e.g., lysine, isoleucine, valine, etc) and their breakdown products, suggesting that proteolysis and/or amino acid catabolism are particularly affected at the whole-body level, followed by reperfusion. On the other hand, top 50 metabolites that contribute to component 2 were enriched in blood samples collected after ischemia without reperfusion (light blue in **Fig. 1C**). Many of these metabolites were associated with carbohydrates (e.g., arabinose, allose, glucose-6-phosphate, etc), suggesting that ischemia and subsequent hypoxia or nutrient shortage in the heart influence carbohydrate metabolism at the whole-body level.

Previously, Goetzman et al. conducted a similar metabolomics study in the systemic venous blood from the jugular vein of pigs following I/R and found perturbations in fatty acids, TCA cycle metabolites, and amino acids^17^. When we examined these metabolites in the systemic arterial blood, we found consistent changes (**Fig. 1D-G**). For example, as reported by Goetzman et al., we also observed significant decreases in multiple long-chain polyunsaturated fatty acids (PUFAs) following ischemia compared to baseline conditions (**Fig. 1G**). These decreases were generally maintained but became more pronounced after reperfusion. In contrast, ischemia tended to increase the circulating levels of several glycolysis (**Fig. 1D**) and TCA cycle intermediates (**Fig. 1E**) at 60 minutes of ischemia. Pyruvate and lactate levels seemed to be further increased after reperfusion, whereas glucose and TCA cycle metabolites’ levels tended to recover to basal levels. Together, these shifts likely reflect a metabolic transition from fatty acid oxidation to anaerobic glycolysis, the hallmark of ischemia^18,19^. Finally, we observed the impact of ischemia on the circulating levels of specific N-acetylated amino acids, consistent with findings from Goetzman et al. (**Fig. 1F**). Thus, our arterial blood metabolite profiles confirmed these previously reported findings in the systemic venous blood. Our results motivated us to determine whether the origin of the altered circulating metabolites during I/R is the heart.

### Ischemia paradoxically enhances cardiac uptake of circulating fatty acids

To investigate how I/R influences the metabolic activity of the heart, we employed heart-specific AV metabolomics methods that we and others have established^8–15^ (see Methods). Specifically, comparing the metabolite abundances between the arterial blood (A) and the coronary sinus (CS) enables us to infer net cardiac uptake or release of individual metabolites (**Fig. 1A**). This analysis provides unique insights into the cardiac nutrient utilization and metabolite production in response to I/R, information hardly captured from snapshot metabolite levels.

PLS-DA analysis of AV metabolomics data revealed discrete clustering by each time point (**Fig. 2A**), indicating that cardiac metabolism changes distinctly over time during I/R. We first examined whether these changes in the metabolism of the heart at least in part contribute to the alterations in the systemic blood that we observed (**Fig. 1C**). Analysis of the top 50 discriminating metabolites in components 1 and 2 for AV metabolomics data (**Fig. 2B**) identified select metabolites whose levels were also changed in the systemic blood during I/R (**Fig. 1C**). For example, several amino acids (e.g., tryptophan, phenhlyalanine, leucine, etc) exhibited altered uptake/release by the heart after reperfusion (**Fig. 2B**), while their levels were changed in the systemic blood as well (**Fig. 1C**). On the other hand, we identified metabolites whose uptake/release by the hearts were changed in response to I/R but their systemic blood levels were unchanged. They were nucleotides and nucleotide derivatives (e.g., cytidine, deoxyuridine) and microbiome-related metabolites (e.g., hydroxyhippuric acid, indole-3-pyruvate, indoxyl sulfate), suggesting that they originated from non-cardiac tissues through the indirect effects of cardiac I/R.

**Figure 2.**
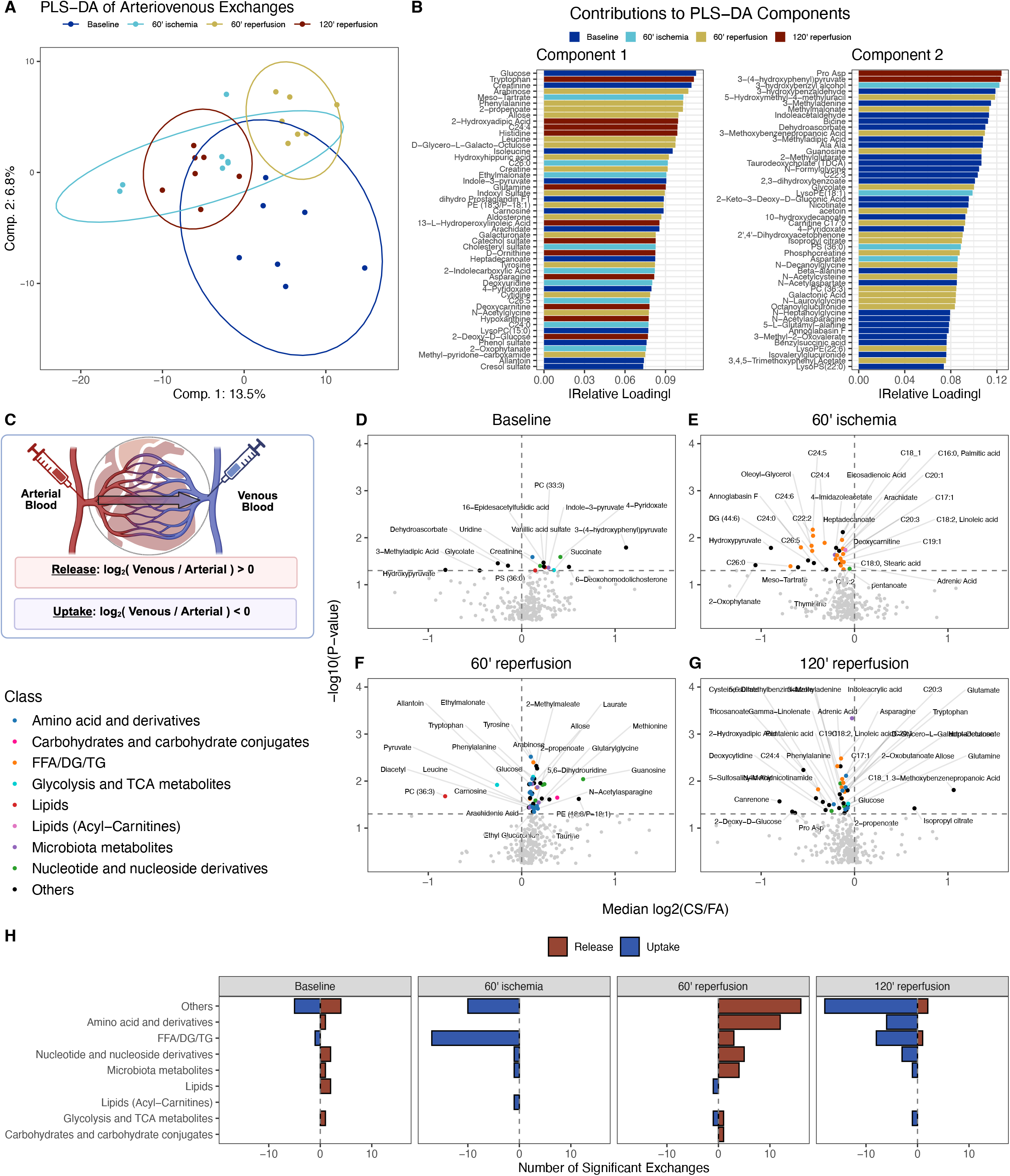
Distinct metabolite uptake and release by the heart during ischemia and reperfusion time course. **A,** PLS-DA score plot showing the separation of arteriovenous (AV) metabolite exchange profiles across four ischemia and reperfusion timepoints. **B,** Top metabolites contributing strongly to the components 1 and 2. **C,** Schematic of metabolite gradient measurements to identify net release or uptake by the heart. **D-G:** Volcano plots for each time point illustrating significant metabolite exchanges through the myocardium. **H:** Bar plot illustrating the number of metabolites showing significant net uptake (leftward bars) or release (rightward bars) at each time point during the ischemia-reperfusion protocol. Metabolites are grouped on the y-axis by biochemical class (e.g., amino acids, lipids, organic acids), and the x-axis shows the count of significantly altered metabolites within each group.

To gain more specific insights into how I/R influences the metabolic activities of the heart, we generated volcano plots of AV metabolomics data for each time point during I/R and color-coded different categories of metabolites (**Fig. 2D-G**). We found a total of 137 metabolites that showed significant uptake/release by the heart across the time series. Unexpectedly, ischemic hearts displayed a marked uptake in circulating non-esterified long-chain fatty acids (LCFA) (**Fig. 2E and Supplementary Fig.1**), compared to the baseline (**Fig. 2D**). This observation is paradoxical, as ischemia is known to increase myocardial utilization of glucose rather than LCFAs as fuel^18^. Therefore, the observed increase in LCFA uptake may reflect a compensatory response in non-ischemic myocardial regions, where enhanced fatty acid oxidation maintains contractile function^18^. Conversely, within ischemic zones, LCFAs and their acylcarnitine derivatives accumulate within minutes of ischemia onset due to altered mitochondrial transport dynamics and inhibited fatty acid oxidation. Indeed, ischemia increases carnitine palmitoyl transferase 1 (CPT1) activity while impairing CPT2, promoting conversion of fatty acids to acylcarnitines that accumulate in the mitochondrial intermembrane space^20,21^. While our data do not allow spatial resolution of these processes, the observed lipid uptake is consistent with early shifts in fatty acid handling that have been documented in ischemic heart tissue.

### Reperfusion changes cardiac fatty acid trafficking in a time-dependent manner

We next examined the effect of reperfusion on cardiac metabolism. Intriguingly, at 60 minutes of reperfusion, the trend of LCFA uptake after ischemia markedly reversed (**Fig. 2F**). For example, the heart released arachidonic acid after reperfusion. Arachidonic acid release likely reflects the increased activity of phospholipase A_2_ (PLA_2_), which produces arachidonic acid by cleaving phospholipids during I/R tissue injury^22^. The heart also released creatine, creatinine, several amino acids, nucleotides and their derivatives (**Fig. 2E**). While creatine kinase release is a well-established marker of cardiomyocyte necrosis during I/R^23,24^, our observation of creatine release may indicate the disruption of intracellular phosphocreatine pool. Cardiomyocytes import creatine via SLC6A8 and store it as phosphocreatine for rapid ATP regeneration^25^. During ischemia, phosphocreatine is quickly depleted to sustain contractile function, which may permit efflux of creatine and its spontaneously formed byproduct, creatinine^25^. Thus, their appearance in coronary sinus effluent likely reflects depleted phosphocreatine pools. Surprisingly, by 120 minutes of reperfusion, we observed increased uptake of LCFAs (**Fig. 2G**), suggesting that the heart has partially recovered from ischemia and re-initiated the oxidation of circulating fatty acids, a process critical for myocardial repair following ischemia^18^. Together, these results, further summarized in **Figure 2H**, reveal time-dependent, dynamic metabolic rewiring of the heart in response to I/R.

### Reperfusion alters cardiac amino acid release in a time-dependent manner

Our comparisons between ischemic and reperfusion timepoints revealed additional patterns of metabolic changes of the heart (**Fig. 3**). In particular, when we compared ischemia with 60 minutes of reperfusion, we identified more unexpected instances of significant metabolite uptake/release by the heart (using one-sample *t*-test), For instance, 60 minutes of reperfusion was marked by significant cardiac release of branched-chain amino acids (BCAAs) (**Fig. 3A**). Intriguingly, similar to LCFAs (**Fig. 3B**), BCAAs also showed reversed patterns by 120 minutes of reperfusion, suggesting the partial recovery of the heart. Importantly, elevated plasma levels of BCAAs have been reported in patients with heart failure, attributed to impaired BCAA oxidation^18^. Previous studies have found a complex role of amino acid metabolism in I/R injury: essential amino acid supplementation reduces injury severity in Langendorff-perfused rat hearts^26^, while mice deficient in BCAA catabolism accumulate BCAAs, leading to suppressed glucose oxidation and increased vulnerability to cardiac injury^27,28^. Notably, BCAA catabolism gene expression is downregulated in both mouse and human failing hearts^28^, which may further contribute to the cardiac release of BCAAs and its elevated levels in blood.

**Figure 3.**
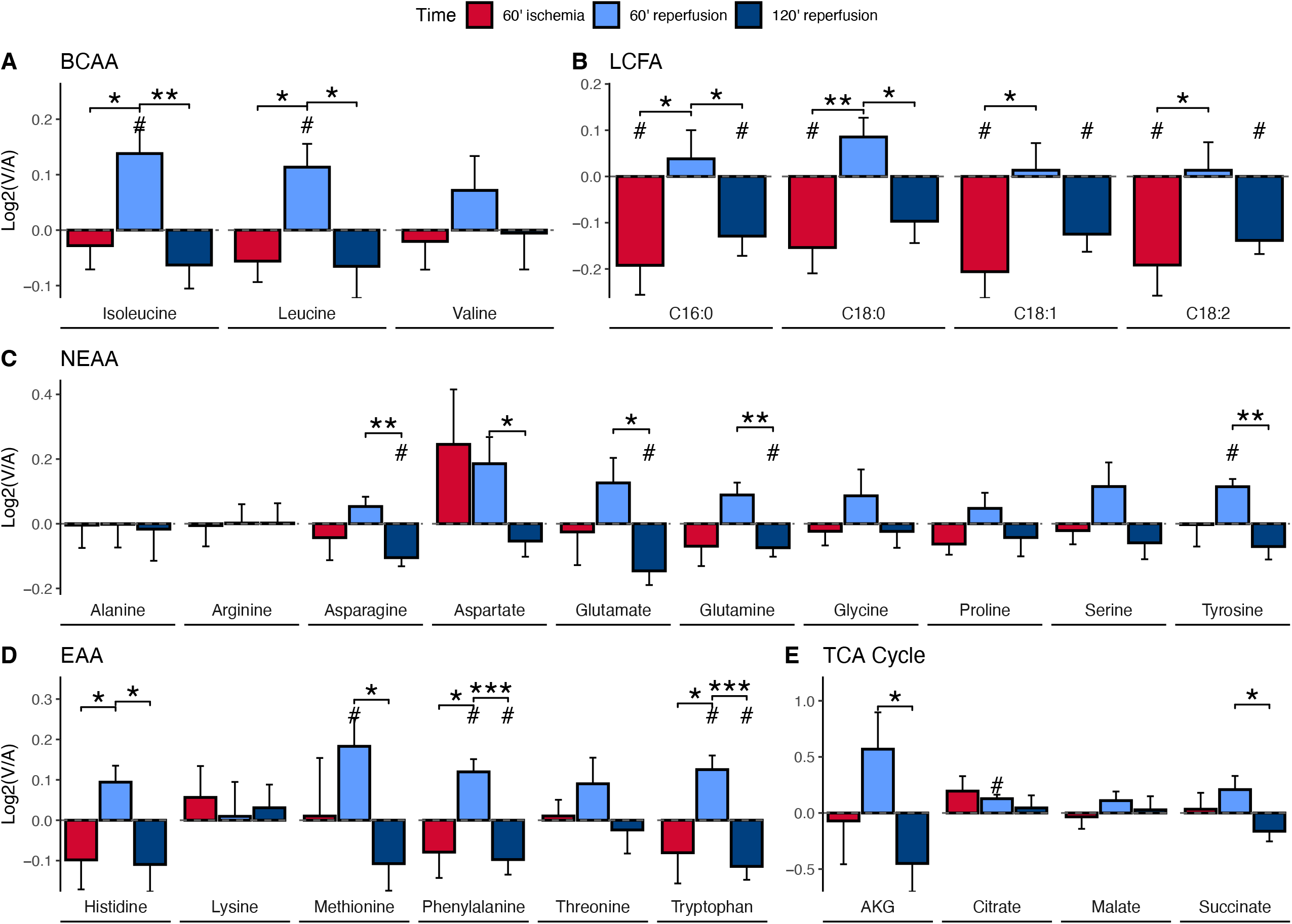
Ischemic hearts take up circulating fatty acids, whereas reperfused hearts at 60 min, but not 120 min, release amino acids and TCA cycle intermediates. A-E: Bar graphs display log2-transformed venous-to-arterial (V/A) ratios for selected metabolites at 60 minutes and 120 minutes post-reperfusion. Positive values (log2[V/A] > 0) indicate net release from the heart, while negative values (log2[V/A] < 0) indicate net uptake. Significant uptake or release events at each timepoint are denoted by # (p < 0.05). Statistically significant changes in flux between timepoints are indicated by *p<0.05, **p<0.01, ***p<0.001.

Compared to BCAAs, other amino acids were less studied in the context of heart diseases. In our dataset, we observed several essential amino acids were also released by the heart after 60 minutes of reperfusion (**Fig. 3D**). Given that essential amino acids were only derived from diet or protein degradation, this finding may reflect active protein breakdown induced by tissue injury after reperfusion following ischemia. Non-essential amino acids were also released by the heart after 60 minutes of reperfusion (**Fig. 3C**), further supporting enhanced proteolysis in this condition. Consistent with this idea, we have reported the substantial release of amino acids from the failing human hearts^15^. In the same study, we have also reported the significant release of TCA cycle intermediates suggestive of anaplerotic stress. Consistently, except for citrate, several TCA cycle metabolites exhibited a trend of release after I/R (**Fig. 3E**).

### Metabolic shifts in transcriptome during I/R may contribute to flux changes

To gain insights into the mechanisms that underly the metabolic flux changes of the heart during I/R, we surveyed the literature for transcriptomic datasets from I/R experiments. Among numerous published datasets, we identified one potentially relevant experiment with timepoints similar to our study: Li et al. analyzed gene expression in a Langendorff-perfused heart model subjected to 20 minutes of ischemia followed by 20 minutes of reperfusion, using RNA-sequencing^29^. Although this model reflects the whole heart ischemia whereas our model induces ischemia in spatially restricted regions of the heart, we reasoned that this dataset would still provide mechanistic insights into our observed flux changes. Our re-analysis of the raw data from Li et al. revealed differentially expressed genes (DEGs) between IR and control hearts (continuous perfusion). The most notable genes were inflammation-associated genes and hypoxia-responsive genes, including multiple heat shock proteins (e.g., Hspa1a, Dnaja1, Bag3), which were highly upregulated after I/R (**Fig. 4A,B**). Heat shock proteins are known to protect myocardial tissue during IR injury by mitigating mitochondrial damage, supporting proteostasis, and preventing excessive mitophagy to sustain ATP production^30–34^. Their induction likely reflects widespread proteotoxic stress and activation of protein quality control mechanisms, including proteasomal and lysosomal degradation of damaged proteins, which may contribute to increased amino acid liberation.

**Figure 4.**
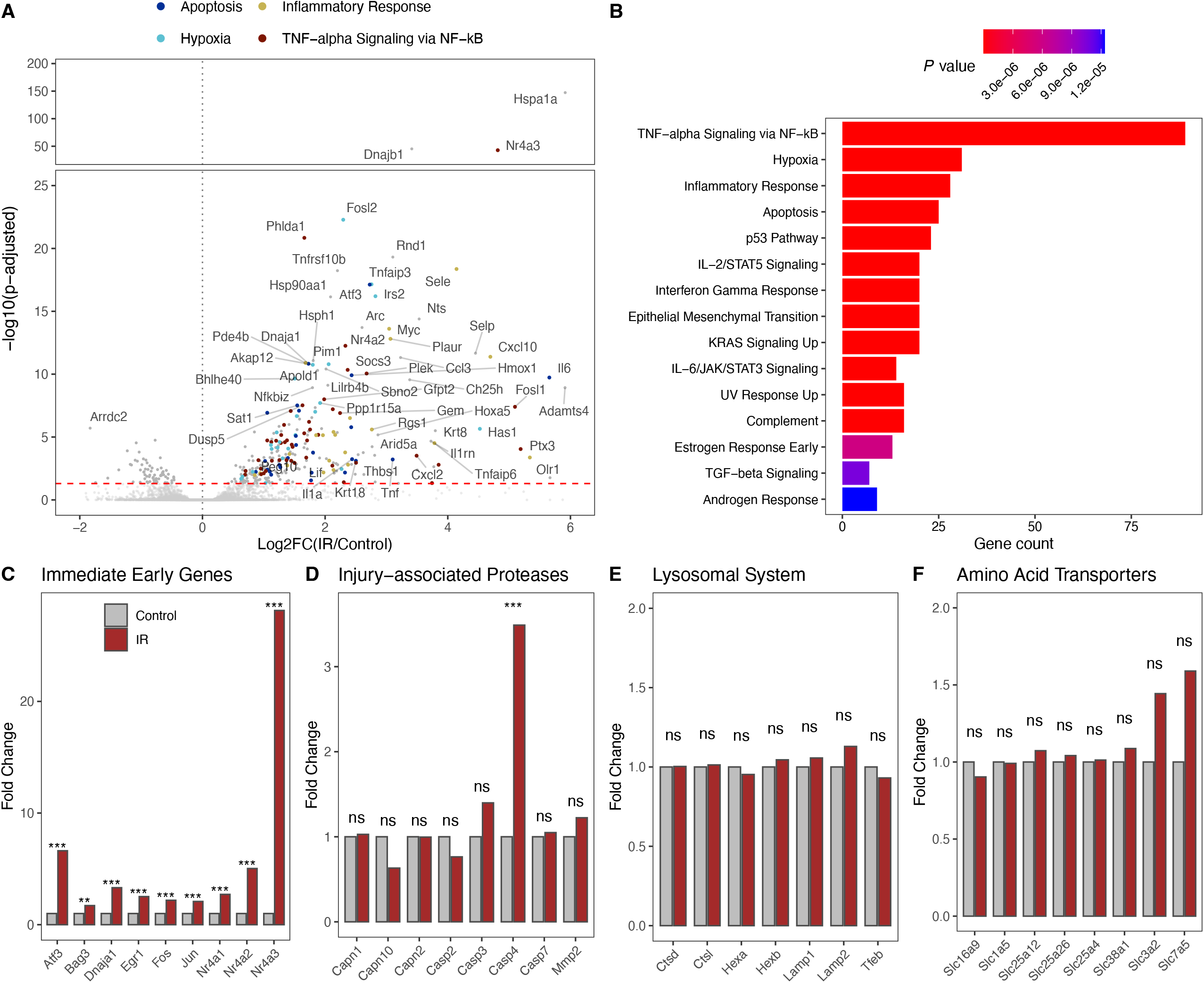
Transcriptomic alternations of the ex vivo perfused mouse hearts after I/R injury. **A,** Volcano plot depicting differential expression of metabolism-related genes 20 min after I/R injury compared to the baseline. **B,** Top biological pathways affected by I/R injury. **C-F,** Bar graphs showing fold changes in the expression of genes related to the indicated pathways. **p<0.01, ***p<0.001 by Student’s t-test.

In addition, members of the NR4A nuclear receptor family (*Nr4a1, Nr4a2*, and *Nr4a3*) were significantly upregulated by I/R (**Fig. 4A, C**). These transcription factors have been implicated in the regulation of metabolic programs, including amino acid metabolism^35^. *NR4A2* expression correlates with MI severity in humans^36^, and *Nr4a1* has been shown to regulate reticulophagy and intracellular amino acid homeostasis, with *Nr4a1* deficiency resulting in reduced amino acid levels^35,37,38^. Consitent with these transcriptomic changes, in our porcine I/R model, we observed net release of both essential and nonessential amino acids from the heart into circulation at 60 minutes post-reperfusion (**Fig. 3**). By 120 minutes, this pattern shifted toward net uptake. Together, these complementary observations from Langendorff-perfused and *in vivo* I/R models suggest a coordinated reperfusion response, in which early activation of proteostatic pathways contributes to protein breakdown and amino acid release, followed by a later shift toward metabolic recovery and anabolic reuptake.

Finally, to investigate whether transcriptional regulation at least in part contributes to the changes in amino acid efflux after I/R, we examined the expression of amino acid transporters, lysosomal genes, and proteases. However, most amino acid transporters were not significantly upregulated in IR hearts, except for Slc3a2 and Slc7a5, which exhibited a modest (∼1.5-fold) trend toward increased expression (**Fig. 4F**). This suggests that the acute amino acid release in vivo is primarily driven by concentration gradients or post-transcriptional mechanisms rather than transcriptional upregulation of transport machinery. Similarly, lysosomal genes showed no significant changes or clear trends (**Fig. 4E**). Interestingly, among the proteases we checked—including calpains, caspases, and metalloproteases-Caspase 4 was significantly upregulated (∼3.5-fold increase, padj < ***). (**Fig. 4D**). Future studies will be necessary to determine the importance of Caspase 4 in I/R hearts.

Collectively, these findings suggest that the early amino acid release observed at 60 minutes post-reperfusion likely reflects acute cellular injury, with initial inflammatory responses being activated to contain the damage. In contrast, transcriptional programs governing lysosomal degradation and proteolytic clearance may be delayed, likely becoming prominent at later reperfusion timepoints.

## Discussion

In this study, we employed heart-specific AV metabolomics in the highly relevant porcine model and revealed dynamic shifts in myocardial substrate utilization during acute I/R. By simultaneously sampling arterial inflow and coronary sinus outflow, we quantified how the ischemic anterior myocardium perturbs whole-heart metabolism. Key findings include a transient suppression of long-chain fatty acid uptake immediately after reperfusion and significant alterations in amino acid handling. These results provide insights into the metabolic distress and potential maladaptive responses of the post-ischemic heart.

### Alterations in Myocardial Energy Metabolism During IR

Consistent with the pathophysiology of myocardial ischemia, we observed a shift toward anaerobic carbohydrate metabolism. It is known that during the ischemic interval (60 min occlusion in our model), the myocardium largely loses oxidative capacity, leading to accelerated glycolysis and lactate accumulation^17^. This lactate buildup in cardiomyocytes is likely due to increased production rather than the uptake of circulating lactate because we do not observe any significant change in lactate uptake/release by the heart during I/R despite increased circulating levels^39^. This sustained glycolytic flux likely reflects both washout of lactate that built up in the ischemic anterior wall and ongoing anaerobic glycolysis in regions with compromised perfusion.

In terms of TCA cycle, succinate is particularly known to accumulate during ischemia and then undergo rapid oxidation on reperfusion in mice, driving a burst of reactive oxygen species (ROS) generation via reverse electron transport in mitochondria^40^. We observed transient spike of succinate in coronary sinus after 60 minutes of reperfusion (**Fig. 3H**), which may corroborate the concept suggested from the mouse study that ischemic succinate accumulation contributes to reperfusion injury by fueling oxidative stress^40^. Our data in a large-animal model thus underscores the translational relevance of targeting succinate metabolism to mitigate IR injury.

Another important metabolic rewiring of the heart that undergoes I/R injury is the disruption of myocardial lipid utilization. Early reperfusion was characterized by an acute impairment of LCFA uptake across the heart. Under normal aerobic conditions, the healthy myocardium derives the majority of its ATP from β-oxidation of fatty acids^17^, extracting substantial amounts of circulating free fatty acids (FFAs) to fuel oxidative phosphorylation. In our IR model, however, the net extraction of FFAs was significantly blunted immediately upon reperfusion. This suggests that the post-ischemic myocardium cannot effectively oxidize fatty acids in the first minutes of reflow, likely due to mitochondrial dysfunction and accumulation of inhibitory fatty-acid intermediates.

Indeed, it is well known that reperfusion after ischemia is accompanied by a surge in FFA supply (from adipose lipolysis and catecholamine stimulation) and a heavy reliance on fatty acid oxidation, which paradoxically can exacerbate myocardial injury^21^. Excess FFA oxidation in the context of ischemia leads to inefficient oxygen use and generates harmful byproducts that stress the heart^41^. One well-documented consequence is the accumulation of long-chain acylcarnitines – partial β-oxidation intermediates that accumulate when fatty acid catabolism is incomplete. These toxic intermediates rapidly build up in ischemic myocardium and are known to disrupt mitochondrial membranes and ionic homeostasis^42^. Although we did not directly measure the accumulation of acylcarnitines within the cardiac tissues, the suppressed FFA uptake and altered organic acid profiles we observed in early reperfusion are consistent with a scenario of incomplete fatty acid oxidation. In other words, the ischemic myocardium initially cannot fully utilize the flood of FFAs upon reperfusion, leading to a mismatch between fatty acid supply and oxidation. This interpretation agrees with prior studies showing that long-chain acylcarnitines increase within minutes of ischemia onset^42^ and likely persist until mitochondrial β-oxidation gradually recovers.

By the late reperfusion phase in our study (120 minutes of reperfusion), myocardial lipid metabolism began to normalize partially. For example, coronary sinus FFA levels in late reperfusion indicated a restoration of net fatty acid extraction compared to early reperfusion, suggesting that surviving myocardium slowly regained the capacity to oxidize fatty acids as perfusion and oxygenation stabilized. This recovery of FFA utilization implies a degree of metabolic flexibility returning to the heart at later time points. Prior work has established that high FFA levels and oxidation during reperfusion can decrease cardiac efficiency and contribute to contractile dysfunction^21^. In pigs and humans alike, elevated plasma FFAs are associated with reduced mechanical efficiency of the left ventricle^43^. Our findings reinforce this concept by demonstrating the heart’s struggle to utilize FFAs immediately after ischemia. The partial recovery achieved through late reperfusion, while encouraging, still suggests a lingering metabolic inflexibility that may affect functional recovery.

Importantly, these metabolic disturbances in substrate utilization lend support to metabolic therapies proposed for myocardial IR injury. The preference for carbohydrate metabolism (glycolysis) over fatty acid oxidation during stress is beneficial at times, as glucose oxidation yields more ATP per oxygen and produces less ROS than fatty acid oxidation^41^. Indeed, interventions that acutely shift myocardial substrate use toward glucose and away from fatty acids have shown cardioprotective effects. For example, administering insulin or dichloroacetate to increase pyruvate oxidation, or using partial inhibitors of fatty acid oxidation, can decrease infarct size and improve post-ischemic cardiac function in animal models^44^. They may imply that the injured heart is initially “forced” into a carbohydrate-dependent state – a condition that therapeutic metabolic modulation attempts to mimic and prolong. Thus, the metabolomic flux profile observed here not only aligns with the known metabolic benefits of glucose-centric therapies but also offers potential biomarkers to gauge the efficacy of such interventions in real-time.

### Alterations in Amino Acid Handling and Anaplerosis

We found that IR altered the heart’s handling of BCAAs. During early reperfusion, there were transient changes in the myocardial uptake/release of BCAAs, suggesting a disruption of normal BCAA catabolism. Under physiological conditions, the heart can oxidize BCAAs as an auxiliary energy source, and indeed evidence shows that myocardium metabolizes BCAAs even at rest^45^. In our study, however, the data indicate an initial accumulation of BCAAs in the coronary sinus (relative to arterial levels) shortly after reperfusion, which suggests that the heart temporarily becomes a net source of BCAAs. One interpretation is that acute ischemia impairs the activity of BCAA dehydrogenases in mitochondria, causing a bottleneck in BCAA utilization and a buildup of BCAAs. This scenario is plausible given that mitochondrial function was compromised early after reperfusion, and similar enzyme inhibition has been noted in heart failure models. The clinical relevance of BCAA perturbation is supported by reports that BCAA levels rise in infarcted myocardium and may contribute to adverse outcomes. For instance, BCAAs have been found to be markedly elevated in cardiac tissue following MI^46^, and excessive accumulation of BCAAs (or their acylcarnitine derivatives) can interfere with glucose oxidation and exacerbate ischemic injury. Conversely, lowering circulating BCAA levels has been shown to mitigate post-MI cardiac dysfunction in experimental studies^46^. Thus, our observation of transient BCAA metabolic disturbance provides a potential link to these findings, suggesting that a failure to adequately oxidize BCAAs in the acute phase of reperfusion may be another facet of metabolic inflexibility that hinders cardiac recovery. By late reperfusion in our model, BCAA gradients began to normalize, indicating that as mitochondrial function improved, the heart may have resumed more normal catabolism of these amino acids. Still, the early imbalance in BCAA handling could have downstream effects on myocardial signaling and remodeling that merit further investigation.

Another insight from our amino acid data is evidence of proteolysis and cellular injury reflected in the coronary sinus amino acid profile. Specifically, we noted increases in amino acids that the myocardium does not significantly catabolize, such as phenylalanine and other aromatic amino acids. The heart cannot oxidize or synthesize phenylalanine; thus any net release of phenylalanine from the heart is a strong indicator of protein breakdown or cell rupture releasing intracellular contents^47^. An elevated trans-cardiac gradient of phenylalanine in our I/R pigs would be consistent with myocardial cell damage in the ischemic region, leading to leakage of free amino acids from the injured cells. Prior literature supports the use of phenylalanine as a marker of net protein loss from the heart^47^. Although we did not directly measure cardiac tissue protein turnover, the circulating evidence of non-metabolizable amino acids provides an external readout of structural injury and tissue catabolism occurring due to I/R. This complements our interpretation of the metabolite fluxes: in early reperfusion, the heart is both metabolically reprogramming (shifting fuel use) and suffering some irreversible injury (releasing components of damaged cells). Together, the changes in amino acid flux underscore that I/R triggers not only shifts in energy substrates but also alterations in nitrogen balance and protein metabolism. Our findings put a spotlight on amino acid pathways – an area underappreciated in acute cardiac ischemia – and suggest they play a role in the myocardial response to injury, whether as alternative fuels (glutamate, BCAAs) or as markers of cell damage (phenylalanine). Future studies could build on this by examining how modulating amino acid availability (for example, glutamate supplementation or BCAA oxidation enhancers) affects functional recovery after MI.

### Mechanistic and Clinical Context

Overall, the metabolomic flux signatures observed in this study align with and extend the existing mechanistic framework of myocardial I/R injury. Ischemia forces the heart into a less efficient metabolic state – heavy reliance on anaerobic glycolysis and limited ATP production – while reperfusion, though necessary for survival of the tissue, unleashes metabolic and oxidative stresses as substrates flood back into the heart^21^. Our direct arteriovenous measurements provide an integrated view of how the whole heart copes with these stresses. Prior research has often measured systemic metabolites or tissue biopsies, but measuring coronary sinus effluent allows attribution of metabolite changes specifically to the heart. Numerous studies, dating back decades, have used coronary sinus sampling of individual metabolites (like lactate) to gauge myocardial metabolism and adequacy of reperfusion^39^. We have expanded this approach using a broad metabolomics panel, thereby capturing coordinated changes across multiple pathways simultaneously. This offers a more holistic picture of cardiac metabolism during I/R and confirms that the porcine heart closely resembles human cardiac metabolic responses, reinforcing its value as a translational model. For example, the injurious impact of excess fatty acid oxidation that we confirm in pigs is likewise a feature in humans with ischemic heart disease^21^. The porcine heart’s substrate preferences and responses to ischemia are very similar to those of the human heart^21^, lending confidence that our findings are clinically relevant.

From a mechanistic standpoint, our data highlight the concept of metabolic inflexibility as a contributor to reperfusion injury. Such metabolic inflexibility can create a vicious cycle, as oxidative impairment leads to ROS generation (from succinate and fatty acid overload) and acid accumulation (from lactate), which in turn further damages mitochondria and contractile function. By documenting these processes in vivo, our study provides a framework for testing metabolic interventions. For instance, the pronounced succinate efflux we observed supports the idea of using pharmacological inhibitors of succinate dehydrogenase at reperfusion to limit ROS burst^48^. Likewise, the early derailment of fatty acid oxidation we noted aligns with the beneficial effects seen with acute use of fatty acid oxidation inhibitors or glucose-insulin infusion in reperfusion therapy^49^. In essence, our findings offer metabolic endpoints that therapies could target: reducing the alanine release (by improving pyruvate oxidation), preventing succinate accumulation, normalizing FFA uptake, and enhancing BCAA catabolism could all be strategies to make the post-ischemic heart more “metabolically flexible” and resilient.

### Methodological Considerations and Limitations

Several important considerations must be noted in interpreting our results. First, because only the anterior left ventricular wall was subjected to ischemia in our experimental design, the coronary sinus blood we sampled represents the integrated metabolic output of the entire left ventricle, including both ischemic and non-ischemic regions. The lateral and inferior LV walls were well-perfused throughout and continued normal metabolism. As a result, the metabolite fluxes we measured are diluted averages of the injured and normal myocardial territories. This likely leads us to underestimate the magnitude of metabolic disturbances occurring in the ischemic region. For example, the anterior wall may have been producing lactate and succinate at excessively high rates; however, the unaffected myocardium was still consuming some lactate and oxidizing substrates, thereby tempering the net changes seen in the coronary sinus. Similarly, any release of amino acids from infarcted myocytes could be partially offset by uptake of those amino acids in remote myocardium for metabolism or protein synthesis. This limitation is inherent to global coronary venous measurements. However, it also emphasizes that despite dilution by normal regions, we still detected robust trans-myocardial changes – underscoring how profound the metabolic shifts in the ischemic region must have been to register at the whole-heart level. In future studies, direct sampling of the regional cardiac vein draining the infarct zone (if technically feasible) or imaging approaches might provide even more localized metabolic information. Nevertheless, our approach mirrors the clinical scenario in which overall cardiac metabolic function is assessed from mixed venous blood.

## Conclusions

In summary, this study demonstrates that myocardial I/R fundamentally alters cardiac metabolism in a phase-specific manner, particularly in the domains of lipid and amino acid processing. Using arteriovenous metabolomics analysis, we showed that the post-ischemic heart undergoes a transient metabolic crisis – characterized by impaired fatty acid oxidation, overflow of TCA metabolites, and amino acid release – before partial metabolic recovery occurs hours later. These findings provide an integrative view of cardiac metabolism during I/R and highlight potential metabolic targets for intervention (such as limiting early fatty acid overload, enhancing glucose utilization, and supporting amino acid anaplerosis). Our discussion situates these results within the context of prior knowledge, affirming established concepts while also providing new insights. Ultimately, improving outcomes after MI may require therapies that address this metabolic inflexibility in addition to restoring blood flow. The novel insights from our porcine model lay the necessary groundwork for such metabolic modulation strategies, underscoring the value of in vivo metabolomic flux assessment in uncovering the hidden biochemical sequelae of I/R injury. With careful translation, these metabolic findings could inform better cardioprotective treatments and monitoring protocols in patients with acute MI.

## Methods

### Ischemia and reperfusion surgeries & blood sampling

Male swine weighing 45–50 kg were used in this study (n=7). Anesthesia was induced using a combination of telazol (2.5 mg/kg) and xylazine (5 mg/kg), followed by maintenance with 5% inhaled isoflurane. Animals were fasted overnight and then intubated the following morning with a 9.0 mm endotracheal tube. Core body temperature was maintained at 37.5 ± 0.5°C using a warming blanket. All animals received an initial intravenous bolus of heparin (100 U/kg), followed by supplemental doses of 1000 U administered every hour. 150 mg bolus of amiodarone was given before the procedure. Mechanical ventilation was provided with room air using volume-controlled settings (North American Dräger Narkomed 2B), with a tidal volume of 10 ml/kg. The respiratory rate was adjusted to maintain a PaCO_2_ of 40 mmHg and an arterial oxygen saturation above 95%. Continuous monitoring of ECG and other vital parameters was performed using the SurgiVet Advisor system (V9200, Smiths Medical). Vascular access was obtained in the bilateral femoral arteries with the 6F sheaths placed percutaneously. In addition, an 8F sheath was placed in the left femoral vein. The left main coronary artery was engaged with a 6F HS guide advanced from the right femoral sheath with fluoroscopic guidance. Coronary angiography was performed with the injection of Omnipaque contrast. The coronary guide wire was inserted in the left anterior descending (LAD) artery over which an appropriately sized balloon was advanced distal to the second diagonal branch. The balloon was inflated, occluding the flow. Vessel occlusion was verified with intracoronary (IC) contrast injection, showing no blood flow beyond the balloon. The vessel remained occluded for 60 minutes, after which the balloon was deflated. A brief coronary angiogram was performed to verify the blood flow to the ischemic region. Arterial (from the femoral artery) and venous (from the coronary sinus) blood samples were taken at baseline, 60 minutes following ischemia, 60 minutes following reperfusion, and 120 minutes after reperfusion. Blood was immediately centrifuged at 10,000xg for 10 minutes, and the plasma was stored at -80 °C until further analysis was performed. Following reperfusion, forty milliliters of 40 mM KCl were administered intracoronarily to arrest the heart. The heart was then removed.

### Metabolomics using LC-MS

10 µL of blood was extracted using 300 µL of ice-cold solvent mixture (acetonitrile:methanol:water, 40:40:20, v/v/v). Samples were vortexed and centrifuged at 16,000 × g for 10 minutes at 4°C. A 70 µL aliquot of the resulting supernatant was transferred to LC-MS vials. Metabolites were separated using hydrophilic interaction liquid chromatography (HILIC) on an XBridge BEH Amide column (2.1 mm × 150 mm, 2.5 µm, 130 Å; Waters) maintained at 25°C. The mobile phases were: solvent A (5% acetonitrile in water containing 20 mM ammonium acetate and 20 mM ammonium hydroxide) and solvent B (100% acetonitrile). The flow rate was 150 µL/min, and the gradient was as follows: 0 min, 90% B; 2 min, 90% B; 3 min, 75% B; 7 min, 75% B; 8 min, 70% B; 9 min, 70% B; 10 min, 50% B; 12 min, 50% B; 13 min, 25% B; 14 min, 20% B; 15 min, 20% B; 16 min, 0% B; 20.5 min, 0% B; 21 min, 90% B; 25 min, 90% B. The autosampler was maintained at 4°C, and 3 µL of sample was injected per run. Metabolite detection and quantitation wer performed using a Q Exactive Plus Hybrid Quadrupole-Orbitrap mass spectrometer (Thermo Fisher Scientific) operated in both positive and negative ion modes. Full-scan spectra were acquired over an m/z range of 70–830 at a resolution of 140,000. Data were processed using Compound Discoverer (Thermo Fisher Scientific) and MAVEN (build 682)^50^. To control technical variability in extraction efficiency and injection volume, 15N-valine was added to the extraction solvent as an internal standard. For accurate arteriovenous (AV) ratio determination, arterial and venous serum samples from each pig were processed and analyzed within the same batch. For systemic metabolite concentration measurements, all arterial serum samples were analyzed together to minimize batch effects. To mitigate inter-animal variation, ion intensities were batch-corrected using the ComBat() function from the sva package (v3.46.0) in R (v4.2.1), treating animal identity as a covariate. After correction, Principle component analysis (PCA) of metabolite levels were generated.

### Statistical Analysis

To assess whether each metabolite exhibited significant trafficking (uptake and release), a one-sample two-sided *t*-test was performed to test whether the mean log_2_(CS/A) across biological replicates (*n* = 7) differed from zero. P-values ≤ 0.05 were considered to have statistically significant trafficking. All statistical tests were conducted in R using the function t.test(AV, mu = 0, alternative = “two.sided”) from the stats package (v3.6.2), where AV represents the vector of log_2_(CS/A) values for each metabolite. Detailed implementation is available in our GitHub repository, within the script “01 process ion counts and compute_AV.R”. Comparisons of metabolite trafficking between timepoints (as shown in Figure 3) were conducted using unpaired two-sided t-tests on the AV values (log_2_(CS/A), n = 7 per timepoint). These tests were performed using the t_test() function from the rstatix package (v0.7.2) in R, with the formula t_test(AV ∼ Time) to assess differences in metabolite trafficking between specific timepoints.

### RNA seq analysis and data integration

To investigate gene expression programs that may underlie our observed arteriovenous metabolite fluxes during reperfusion, we reanalyzed publicly available RNA-sequencing data from Li et al. (GEO accession: GSE133796)^29^. Paired-end FASTQ files were downloaded from the Sequence Read Archive (SRA accessions: SRR9641455– SRR9641466) using fastq-dump from the SRA Toolkit (v3.0.0). Reads were pseudo-aligned using Kallisto (v0.51.1) to a transcriptome index generated from Ensembl release 113 *Mus musculus* GRCm39 cDNA^51^. Differential gene expression between ischemia-reperfusion (IR) and control samples (n = 3 per group) was assessed using the DESeq2 package^52^ (v1.38.3) in R, which was used to calculate log2 fold changes and Benjamini-Hochberg adjusted *p*-values. Detailed implementation is available in our GitHub repository

---

**Supplementary Figure 1.**
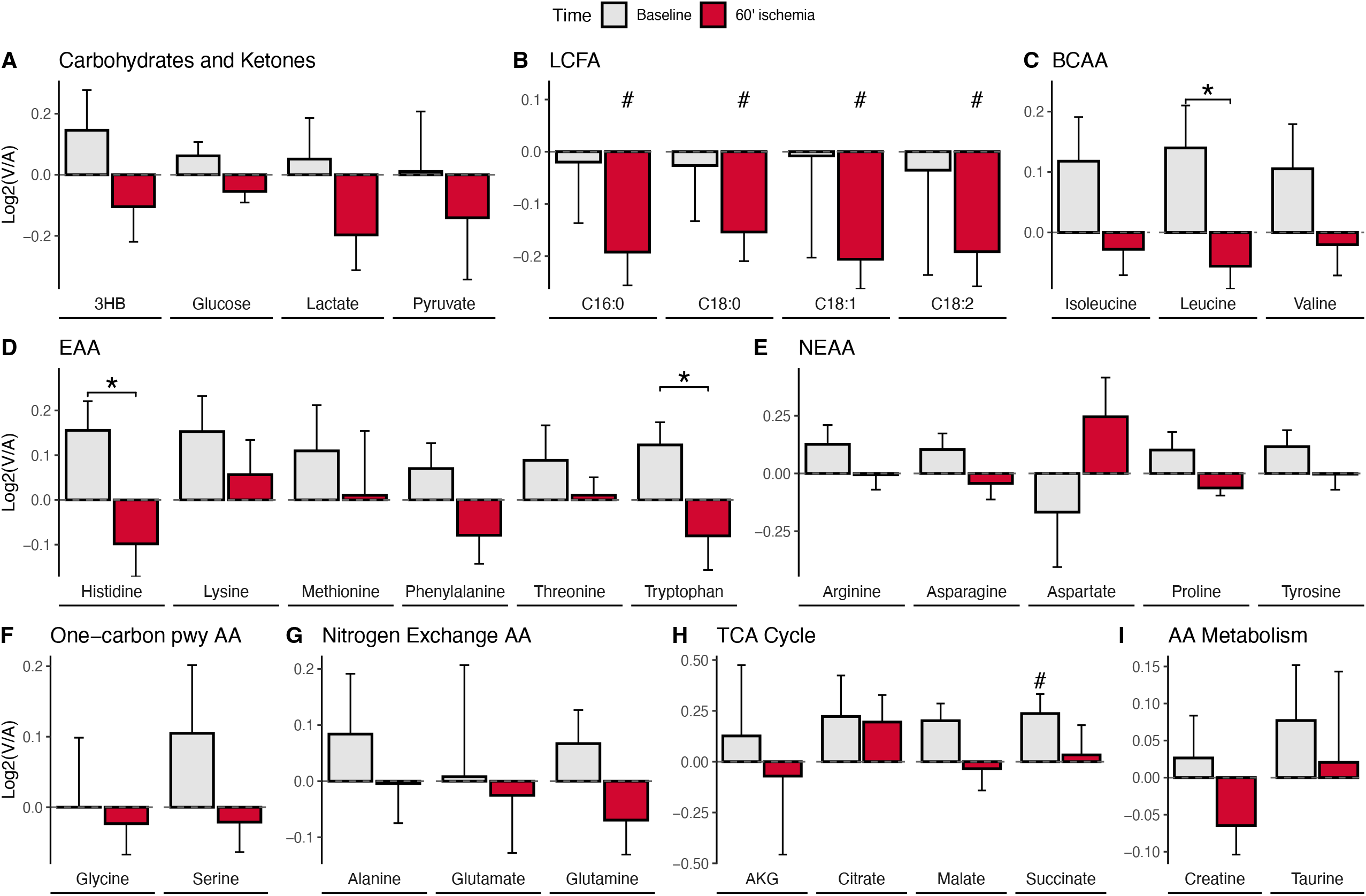
Altered trafficking of major carbon substrates by the hearts in response to ischemia. Bar graphs display log2-transformed venous-to-arterial (V/A) ratios for selected metabolites at 60 minutes and 120 minutes post-reperfusion. Positive values (log2[V/A] > 0) indicate net release from the heart, while negative values (log2[V/A] < 0) indicate net uptake. Significant uptake or release events at each timepoint are denoted by # (p < 0.05). Statistically significant changes in flux between timepoints are indicated by *p<0.05, **p<0.01, ***p<0.001.

